# Multistep metabolic engineering of *Bacillus licheniformis* to improve pulcherriminic acid production

**DOI:** 10.1101/790691

**Authors:** Shiyi Wang, Huan Wang, Dan Zhang, Xiaoyun Li, Jin’ge Zhang, Yangyang Zhan, Dongbo Cai, Xin Ma, Dong Wang, Shouwen Chen

## Abstract

Pulcherriminic acid, a cyclodipeptide, possesses the excellent antibacterial activities by chelating iron ions from environment, however, the low yield has hindered its application. In this study, high-level production of pulcherriminic acid was achieved by the multistep metabolic engineering of *Bacillus licheniformis* DWc9n*. Firstly, leucine (Leu) supply was increased by overexpressing the genes *ilvBHC-leuABCD* and *ilvD* involved in the Leu synthetic pathway and deleting the gene *bkdAB* encoding a branched-chain α-keto acid dehydrogenase. The intracellular Leu content and pulcherriminic acid yield of strain W2 reached 147.4 mg/g DCW and 189.9 mg/L respectively, which corresponded to a 226.7% and 49.1% higher than those of DWc9n*. Secondly, the strain W3 was obtained by overexpressing leucyl-tRNA synthase LeuS in W2, which lead to a 190% improvement compared to DWc9n*, pulcherriminic acid yield increase to 367.7 mg/L. Thirdly, overexpression of the cytochrome synthase gene cluster *yvmC-cypX* further increased the yield of pulcherriminic acid to 507.4 mg/L. Finally, the secretion capability was improved by overexpressing the pulcherriminic acid transporter gene *yvmA*. This resulted in the 556.1 mg/L pulcherriminic acid was produced in W4/pHY-*yvmA*, increased by 340% as compared with DWc9n*, and it is the highest pulcherriminic acid production currently reported. Taken together, this study provided an efficient strategy for enhancing pulcherriminic acid production and might also be applied for high-level production of other cyclodipeptides.

**Importance:** Pulcherriminic acid is a cyclodipeptide that derived from cyclo(L-Leu-L-Leu), which has the same iron chelation group with hydroxamate siderophores. Pulcherriminic acid producing strains can inhibit the growths of various bacteria and plant pathogenic fungi. However, the expected pulcherriminic acid yield was not achieved. This study reports efficient microbial production of pulcherriminic acid in *Bacillus licheniformis* DWc9n* via multistep metabolic engineering strategies. The combination of these interventions resulted in the establishment of a pulcherriminic acid overproducing strain. In combination with bioprocess engineering efforts, pulcherriminic acid was produced at a final yield of 556 mg/L in a shake flask. This is the highest pulcherriminic acid yield ever reported so far using rationally engineered microbial cell factories.

## Introduction

Cyclodipeptides, also known as 2, 5-diketopiperazine or 2, 5-dioxopiperazine, are a class of peptides that are synthesized by cyclodipeptide synthetases (CDPS) (1, 2). At present, a variety of cyclodipeptides with biological activities has been found (3). For example, cyclo(Phe-Pro), cyclo(Tyr-Phe) and cyclo(Leu-Tyr) act as quorum sensing signaling molecules. Cyclo(D-Pro-L-Phe) exhibits antibacterial activity, and cyclo(L-Phe-L-Pro) and cyclo(L-Phe-trans-4-OH-Pro) antagonize fungi (4). Moreover, cyclodipeptides can act as molecular fragments for drug molecular design (3, 5). Therefore, increasing attention has been paid to the search of novel bioactive cyclodipeptides and the construction of recombinant strains with high-level production of cyclodipeptides.

Pulcherriminic acid is an outstanding cyclodipeptide that is derived from cyclo(L-Leu-L-Leu) (6). It is produced by yeasts and *Bacillus* species (7, 8). Iron chelation by pulcherriminic acid to form insoluble pulcherrimin, causing iron depletion in the environment, is thought to mediate antimicrobial activity exhibited by some bacteria and yeasts (9). The producing strains often restrain the growth of various bacteria and pathogenic fungi (7, 10) and have thus been considered as biocontrol and bacteriostatic agents (11–13). Pulcherriminic acid yield was mainly improved by fermentation formula optimization, but it is still at a low level (14).

In recent years, the synthetic pathway of pulcherriminic acid has been extensively studied in *Bacillus* (15), which cleared up the obstacles for enhancing pulcherriminic acid production via metabolic engineering breeding. Firstly, leucine (Leu), the precursor of pulcherriminic acid, is synthesized from glycolytic intermediates, whose formation is catalyzed by IlvBH (acetohydroxyacid synthase), IlvCD (keto-acid reductoisomerase and dihydroxy-acid dehydratase) and LeuABCD (2-isopropylmalate synthase, 3-isopropylmalate dehydrogenase and dehydratase) (16). Leu is then used as the substrate by the leucly-tRNA synthetase LeuS to form leucyl-tRNA (17). The CDPS YvmC then uses two molecules leucyl-tRNA generate cyclo(L-Leu-L-Leu) (cLL), and a cytochrome P450, encoded by *cypX*, oxidizes cLL to form pulcherriminic acid (15). Ultimately, pulcherriminic acid is secreted to the environment by major facilitator superfamily (MFS) transporters (18). Previously, it has been confirmed that the *yvmC*-*cypX* operon is negatively regulated by the transition state regulator AbrB and MarR family regulators YvnA and YvmB (18, 19).

The industrial bacitracin production strain *Bacillus licheniformis* DW2 is generally regarded as safe (GRAS) due to its non-pathogenicity. Previously, an efficient CRISPR/Cas9 nickase mediated genome editing tool was established in *B. licheniformis* DW2 and a bacitracin synthesis-deficient strain *B. licheniformis* DWc9n* was constructed using this toolkit (20). In this work, we aimed to construct an efficient pulcherriminic acid production strain based on the original strain DWc9n*. We employed a multistep metabolic engineering strategy including the strengthening of the carbon flux towards the precursors Leu and leucyl-tRNA and the overexpression of cyclodipeptides synthetase and pulcherriminic acid transportation pathways (Fig. 1). Collectively, this study provided a serviceable strategy as well as a promising *B. licheniformis* for high-level production of pulcherriminic acid.

**Fig. 1.**
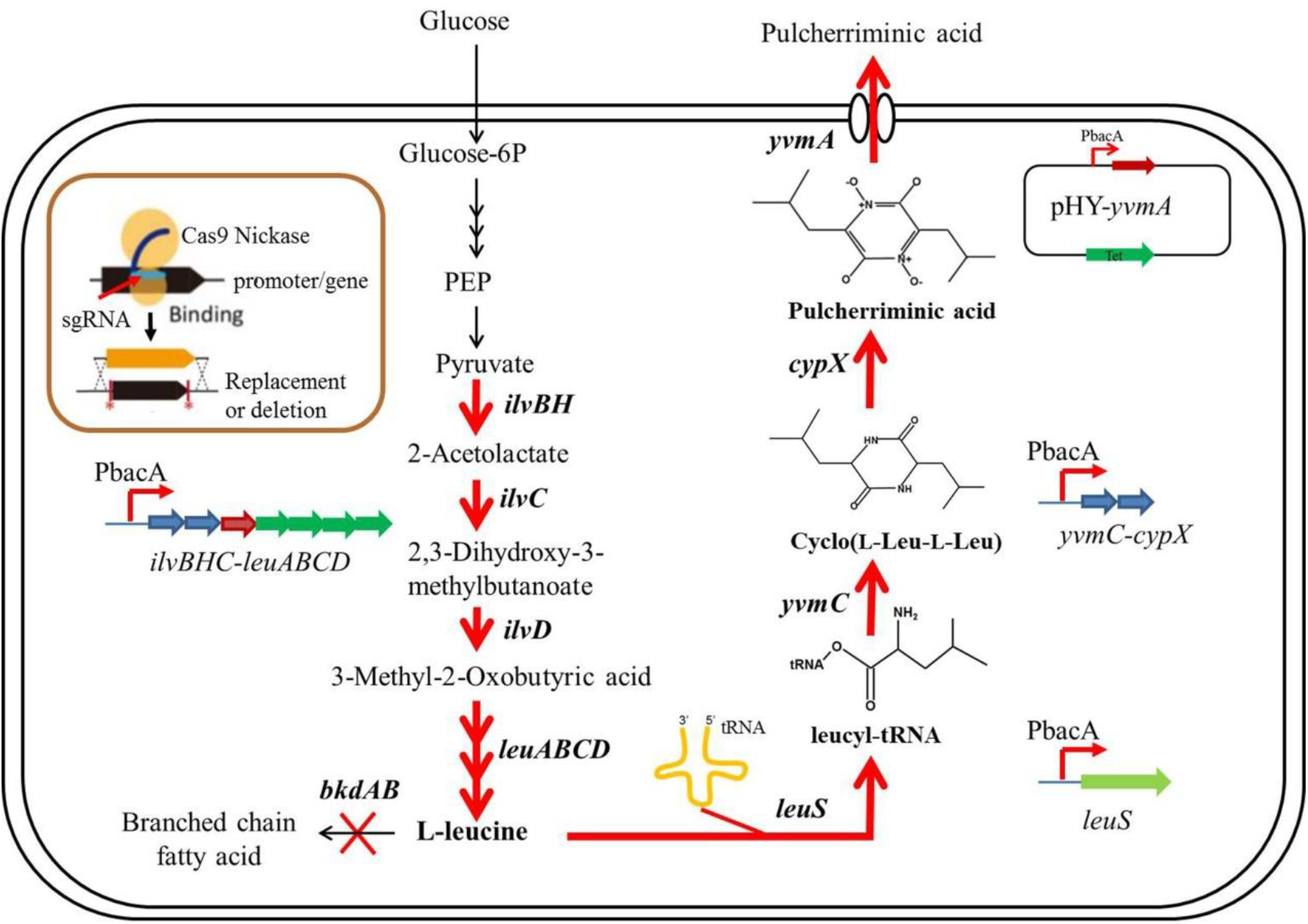
Metabolic engineering of *B. licheniformis* for enhanced production of pulcherriminic acid. Red arrows indicate the overexpressed pathways, multiple arrows represent multi-step reactions, and the red cross indicates a deletion.

## Materials and methods

### Strains, plasmids, and growth conditions

The strains and plasmids used in this study are presented in Table 1. The oligonucleotide primers used in this study are listed in **Table S1**. *Escherichia coli* DH5α was used for vector construction, and *B. licheniformis* DWc9n* was used as the wildtype strain to construct recombinant strains (20). The plasmid pHY300 was used for constructing gene expression vector. Luria Bertani (LB) medium was applied for cultivating *E. coli* and *B. licheniformis*, if necessary, antibiotics (20 μg/L tetracycline, 20 μg/L kanamycin; Sigma-Aldrich) were added to the media. The seed culture was cultivated at 37°C for 12 h, and then transferred into the pulcherriminic acid production medium (g/L: glucose 40, sodium citrate 12, (NH4)_2_SO_4_ 6.20, K_2_HPO_4_·3H_2_O 0.50, MgSO_4_·7H_2_O 0.50, CaCl_2_·2H_2_O 0.20, FeCl_3_·6H_2_O 0.25, and MnSO_4_·H_2_O 0.02, pH 7.50) in 250-mL flasks on a rotary shaker (230 rpm) at 37°C for 48 h.

**Table 1.** The strains and plasmids used in this research.

| Strains and plasmids | Description | Source/reference |
| --- | --- | --- |
| **Strains** |  |  |
| *E. coli* DH5α | *supE44 ΔlacU169 (f 80 lacZΔM15) hsd R17 recA1 gyrA96 thi1 relA1* | TaKaRa Co., Ltd |
| *B. licheniformis* DWc9n* | *B. licheniformis* DW2 derivative with Cas9n integrated expression and *bacABC* operon knocked out | Laboratory stock |
| W1 | DWc9n* derivate, PbacA-*ilvBHC-leuABCD-ilvD* | This study |
| W2 | W1 derivate, *ΔbkdAB* | This study |
| W3 | W2 derivate, PbacA-*leuS* | This study |
| W4 | W3 derivate, PbacA-*yvmC-cypX* | This study |
| W4/pHY300 | W4 with pHY300 | This study |
| W4/pHY-*yvmA* | W4 with pHY-*yvmA* | This study |
| **Plasmids** |  |  |
| pHY300 | *E. coli-Bacillus* shuttle vector; Amp<sup>r</sup> in *E. coli*, Tc<sup>r</sup> in both *E. coli* and *B. licheniformis* | Stored in lab |
| pHY-*yvmA* | pHY300 derivative harboring *yvmA* expression cassette for over expression of *yvmA*, Amp<sup>r</sup>, Tc<sup>r</sup> | This study |
| pGRNA | pHY300+P43(noRBS)+gRNA+TamyL(target *yvmC*) | Laboratory stock |
| pGRNA-P<sub>bacA</sub>(*ilvB*) | pHY300+P43(noRBS)+gRNA(PilvB)+TamyL+LH+PbacA+RH*ilvB* | This study |
| pGRNA-P<sub>bacA</sub>(*ilvD*) | pHY300+P43(noRBS)+gRNA(PilvD)+TamyL+LH+PbacA+RH*ilvD* | This study |
| pGRNA-P<sub>bacA</sub>(*leuS*) | pHY300+P43(noRBS)+gRNA(PleuS)+TamyL+LH+PbacA+RH*leuS* | This study |
| pGRNA-P<sub>bacA</sub>(*yvmC*) | pHY300+P43(noRBS)+gRNA(PyvmC)+TamyL+LH+PbacA+RH*yvmC* | This study |
| pGRNA-*ΔbkdAB* | pHY300+P43(noRBS)+gRNA(*bkdAB*)+TamyL+LH+RH*bkdAB* | This study |

### Construction of promoter replacement strains

The promoter of the bacitracin synthetase cluster P_bacA_ has been proven as a strong promoter in our previously reported research (21). Here, the promoter P_bacA_ was employed to replace the original promoters of *ilvBHC-leuABCD* operon, *ilvD, yvmC-cypX* operon, and *leuS* using the CRISPR-Cas9n toolkit to construct the gene overexpression strains. The construction procedure for the *ilvBHC-leuABCD* cluster promoter replacement strain served as an example. Briefly, the ribosome-binding site (RBS)-free P_43_ promoter coupled with the sgRNA fragment, the P_bacA_ promoter, as well as up and down-stream homology arms of P_ilvBHC-leuABCD_ (0.5 kb) were amplified with the corresponding primers (**Table S1**). Then, these fragments were fused by Splicing Overlap Extension (SOE)-PCR and inserted into pHY300 by using the *Eco*RI and *Xba*I restriction sites; constructing the plasmid named as pGRNA-P_bacA_ (*ilvBHC-leuABCD*).

Then, pGRNA-P_bacA_ (*ilvBHC-leuABCD*) was electro-transformed into DWc9n* and the positive colonies were verified by diagnostic PCR and DNA sequencing. Colonies were cultivated in LB medium without tetracycline at 37°C for several generations to remove the remaining plasmids in positive colonies. Then, the tetracycline-sensitive colonies were verified with diagnostic PCR to confirm the identification of the IlvBHC-leuABCD overexpression strain. Similarly, the *ilvD, yvmC-cypX* operon and *leuS* overexpression strains were obtained by the same method.

### Construction of gene *bkdAB* deletion strain

The *bkdAB* deletion strain of *B. licheniformis* was obtained by using the CRISPR-Cas9n toolkit (20) and the same procedure as for promoter replacement. Briefly, the P_43_ promoter and up and down-stream homology arms of *bkdAB* (0.5 kb) were amplified and the gene deletion plasmid pGRNA-∆*bkdAB* was constructed. Then, pGRNA-∆*bkdAB* was transformed into *B. licheniformis* by electroporation and the *bkdAB* deletion strain was verified by diagnostic PCR and DNA sequencing.

### Construction of *yvmA* overexpression strains

The gene expression vector was constructed according to the previously reported research (22). Briefly, the P_bacA_ promoter, the *yvmA* gene, and the *amyL* terminator from *B. licheniformis* DW2 were amplified and fused by SOE-PCR. The fused fragment was inserted into pHY300 at the restriction sites *Bam*HI and *Xba*I. Named pHY-*yvmA* diagnostic PCR and DNA sequencing confirmed that the YvmA expression vector was constructed successfully. Then pHY-*yvmA* was electro-transferred into *B. licheniformis* to construct YvmA overexpression strain. In addition, pHY300 was transformed into *B. licheniformis* to serve as the control strain.

### Determinations of pulcherriminic acid concentration and cell density

The concentration of pulcherriminic acid was measured according to our previously reported research (14). The purification and standard curve method of pulcherriminic acid was performed according to standard protocols (14). Briefly, the volume of 1 mL culture broth was evenly mixed with 1 mL 2 M NaOH solution to dissolve pulcherriminic acid, centrifuged at 10,000 *g* for 2 min, and the absorbance of supernatant was determined at 410 nm (including intracellular and extracellular pulcherriminic acid). The concentration of pulcherriminic acid was calculated via a standard curve. Meanwhile, the cell density was measured by determining the optical density at 600 nm (OD_600_).

To detect the intracellular pulcherriminic acid, the volume of 1 mL cells was harvested by centrifugation at 10,000 g for 5 min. Then, the cell pellet was washed with distilled water to remove the extracellular pulcherrimin. Afterwards, the cells were disrupted by sonication on ice for 20 min. Cell debris was mixed with 1 mL 2 M NaOH solution to dissolve pulcherriminic acid centrifuged at 10,000 *g* for 2 min, and the absorbance of supernatant was determined at 410 nm.

### Quantitative real-time PCR

*B. licheniformis* cells were collected at 12 h for RNA extraction according to our previous report using an RNA extraction kit (23). The RNA concentration was measured on a NanoDrop 2000 spectrophotometer (Thermo Scientific, USA). The first-strand cDNA was synthesized from 50 ng total RNA using Revert Aid First Strand cDNA Synthesis kits (Thermo, USA). The resulting cDNA was used as the template for qRT-PCR with primers listed in **Table S1** using the iTaqTM Universal SYBR^®^ Green Supermix (BIO-RAD, USA). The *16S rRNA* served as reference gene and all assays were performed in triplicate.

### Determination of the concentrations of intracellular amino acids

The intracellular precursor amino acids were measured using gas chromatography (GC), according to our previously reported method (16). Concentrations were calculated via standard curves made with the corresponding amino acids.

## Results

### Improving precursor Leu supply improved pulcherriminic acid production

Leucyl-tRNA is generated by Leu that derived from pyruvate serves as the sole substrate for pulcherriminic acid synthesis. The adding of Leu to the medium could significantly increase pulcherriminic acid production of DWc9n* strain (Fig. 2A). It indicated that the inadequate Leu synthesis of DW2c9n* may be one of the important factors limiting pulcherriminic acid production.

**Fig. 2.**
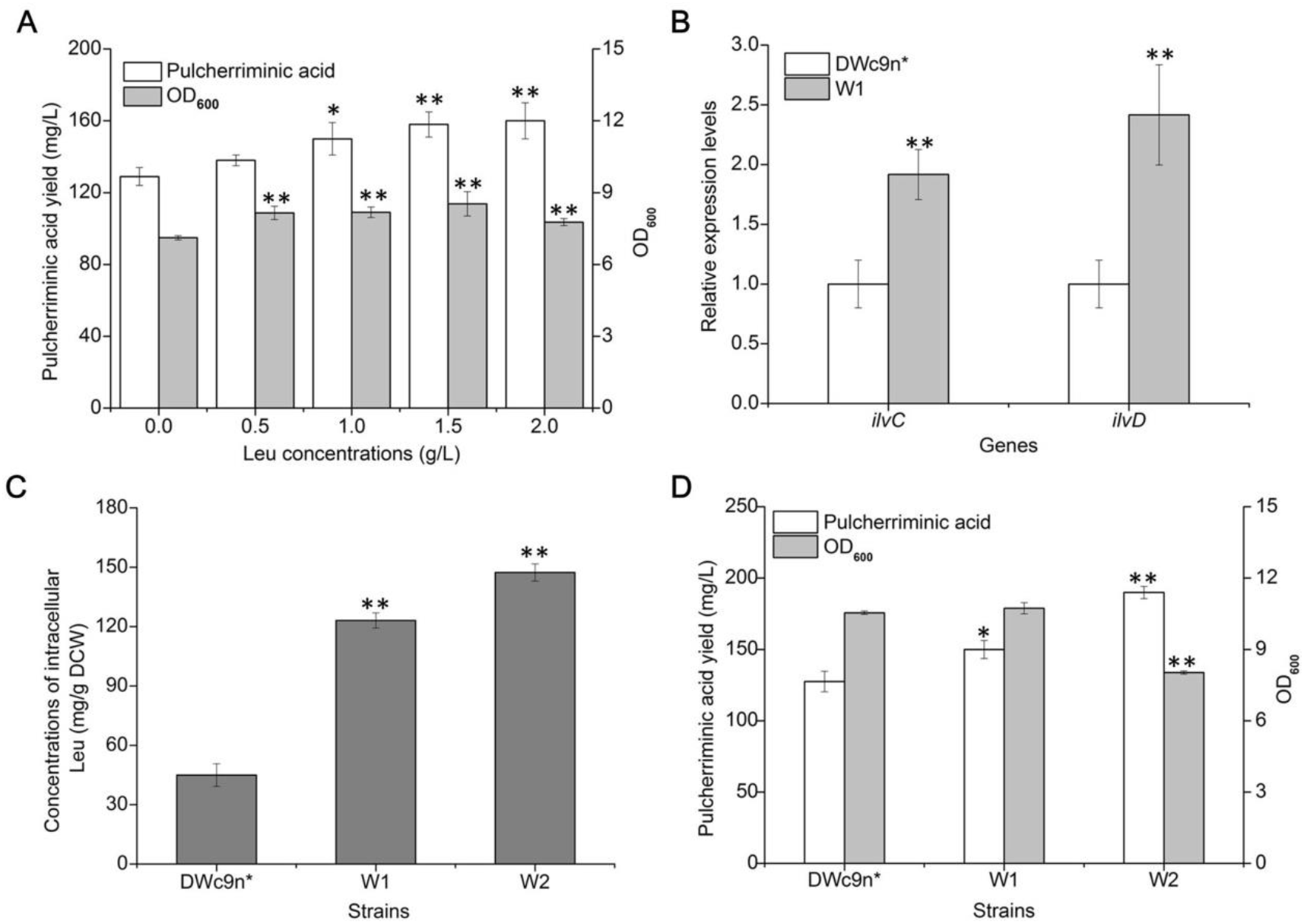
*ilvBHC-leuABCD* and *ilvD* overexpression and *bkdAB* deletion lead to increased pulcherriminic acid production. A: The yield of pulcherriminic acid and OD_600_ under different Leu concentrations in DWc9n*; B: Gene transcriptional levels of the *ilvBHC-leuABCD* operon and *ilvD* at 20 h; C: The concentrations of intracellular Leu at 24 h; D: Pulcherriminic acid yield & OD_600_. All samples were performed in three replicates, and data were presented as the mean ± the standard deviation for each sample point. All data were conducted to analyze the variance at P < 0.05 and P < 0.01, and a *t* test was applied to compare the mean values using the software package Statistica 6.0 ∗, P < 0.05 and ∗∗, P < 0.01 indicated the significance levels between recombinant strains and control.

In order to enhance Leu biosynthesis, the promoters of the *ilvD* gene and the *ilvBHC-leuABCD* operon were replaced by the strong promoter P_bacA_ successively to construct the W1 strain. The influences of promotors replacement on transcription of *ilvD* and *ilvBHC-leuABCD* operon were evaluated by reverse transcription-quantitative PCR (RT-qPCR) at 20 h. The transcription levels of genes in DWc9n* were set to 1. The results showed that transcriptional levels of *ilvD* and *ilvC* were increased by 240% and 190% as compared with DWc9n*. In addition, the intracellular Leu concentration was increased to 123.1 mg/g DCW by 170% as compared to DWc9n* (50.0 mg/g DCW) at 24 h (Fig. 2B&C). Consistent with the increasing Leu concentration, the yield of pulcherriminic acid of W1 reached 149.9 mg/L, increased by 18.1% as compared to DWc9n*(Fig. 2D).

The *bkdAB* gene cluster encodes E1 and E2 components of the branched-chain α-keto acid dehydrogenase complex, which convert branched-chain amino acids (Leu, Ile, Val) into branched-chain α-ketoacyl-CoA starters (BCCSs) and finally be used to the synthesis of branched-chain fatty acids (24). In order to improve the accumulation of intracellular Leu, the *bkdAB* gene was deleted in W1 to obtain strain W2. Based on our results, the concentration of intracellular Leu was 147.4 mg/g DCW in W2, which was 19.7% higher than that of W1 (Fig. 2C). But since branched-chain fatty acids serve as an important component of cell membrane (Bentley et al., 2016), deletion of *bkdAB* was not conducive to cell growth (Wu et al., 2019), and cell density showed 23.8% decrease in W2 as compared to DWc9n*. Certainly, the increase of pulcherriminic acid yield might be another reason for the decrease of cell density. Taken together, deletion of *bkdAB* further increased pulcherriminic acid production and the yield of W2 reached 189.9 mg/L, increased by 26.7% as compared with that of W1 (Fig. 2D).

### Strengthening leucyl-tRNA synthase LeuS expression increased pulcherriminic acid production

CDPS did not activate amino acids but stole the activated amino acids in the form of aa-tRNAs from aa-tRNA synthetases to catalyze the formation of cyclodipeptides (25). Here, overexpression of *leuS* might thus strengthen the conversion of Leu to leucyl-tRNA and further benefit pulcherriminic acid production. The native promoter of *leuS* was replaced by P_bacA_ in the W2 strain to construct the W3 strain. The results showed that the transcriptional level of *leuS* in W3 was 240% higher than that in W2 (Fig. 3A), and the intracellular Leu concentration decreased by 30.2% (Fig. 3B). the pulcherriminic acid yield of W3 reached 367.7 mg/L, increased by 93.6% and 190% compared to W2 and DWc9n* respectively (Fig. 3C). It indicated that overexpression of *leuS* promotes the flow of Leu to the synthesis of pulcherriminic acid.

**Fig. 3.**
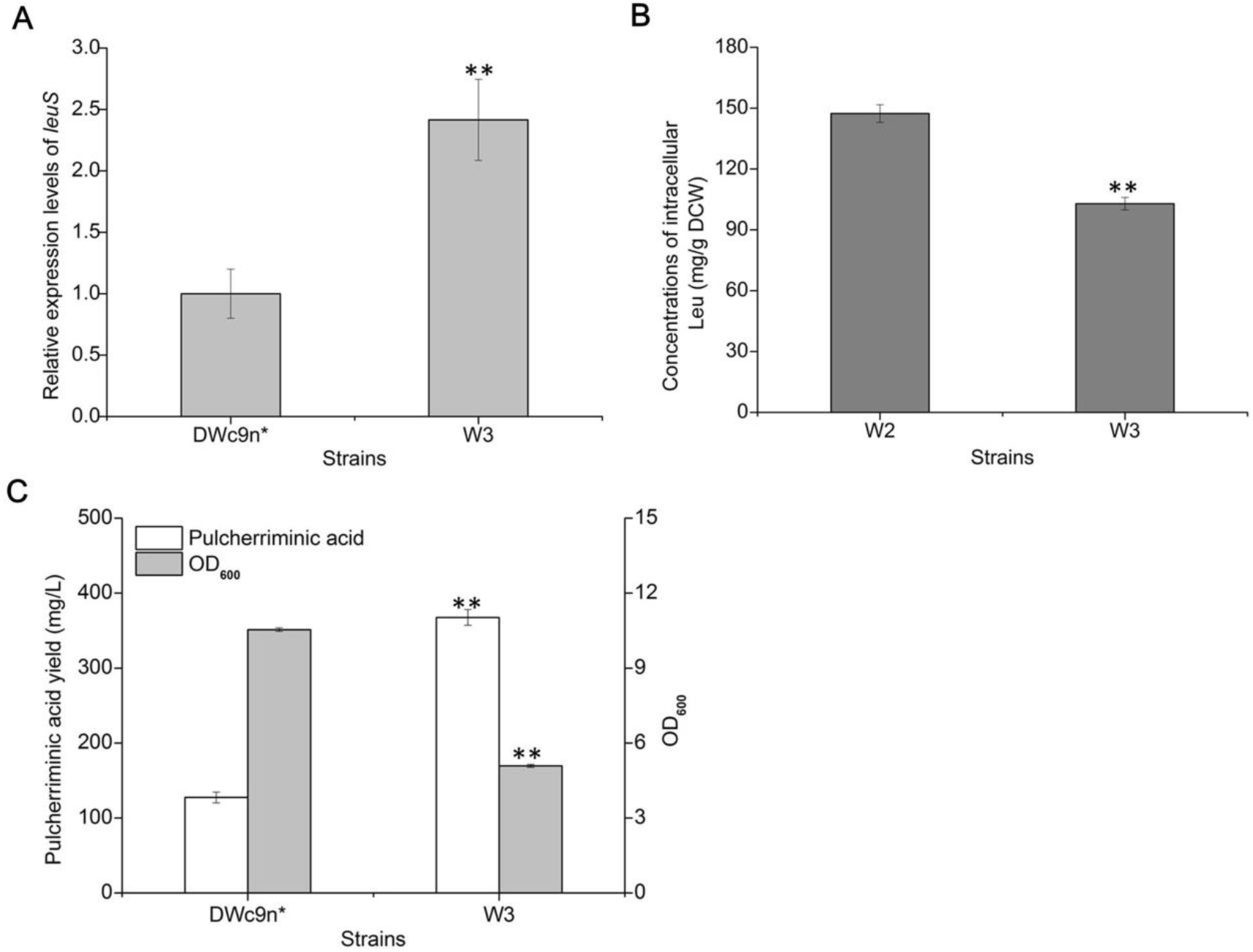
Effect of *leuS* overexpression on pulcherriminic acid production. A: Gene transcriptional levels of the *leuS* at 20 h; B: The concentrations of intracellular Leu at 24 h; C: Pulcherriminic acid yield & OD_600_. All samples were performed in three replicates, and data were presented as the mean ± the standard deviation for each sample point. All data were conducted to analyze the variance at P < 0.05 and P < 0.01, and a *t* test was applied to compare the mean values using the software package Statistica 6.0 ∗, P < 0.05 and ∗∗, P < 0.01 indicated the significance levels between recombinant strains and control.

### Enhancing the expression of the *yvmC*-*cypX* gene cluster promoted pulcherriminic acid production

Although the *leuS* gene was overexpressed to increase precursor leucyl-tRNA formation for pulcherriminic acid synthesis, but the concentration of intracellular Leu in W3 strain was still maintained at a high level. It’s indicating that low expression level of CDPS might also be a bottleneck for pulcherriminic acid production. So, the *yvmC-cypX* overexpression strain was obtained via promoter replacement in the strain W3, and resulted in the recombinant strain W4.

qRT-PCR showed that the transcriptional level of *yvmC* in the W4 strain was 550% higher than in DWc9n* (Fig. 4A). At the same time, the concentration of intracellular Leu in W4 (55.3 mg/g DCW) was decreased by 46.3% as compared with that of W3 (102.9 mg/g DCW) (Fig. 4B). As expected, pulcherriminic acid yield of W4 reached 507.4 mg/L, increased by 300% as compared to DWc9n* (Fig. 4C). These results indicated that the *yvmC*-*cypX* gene cluster was the rate-limiting step for pulcherriminic acid synthesis in W3.

**Fig. 4.**
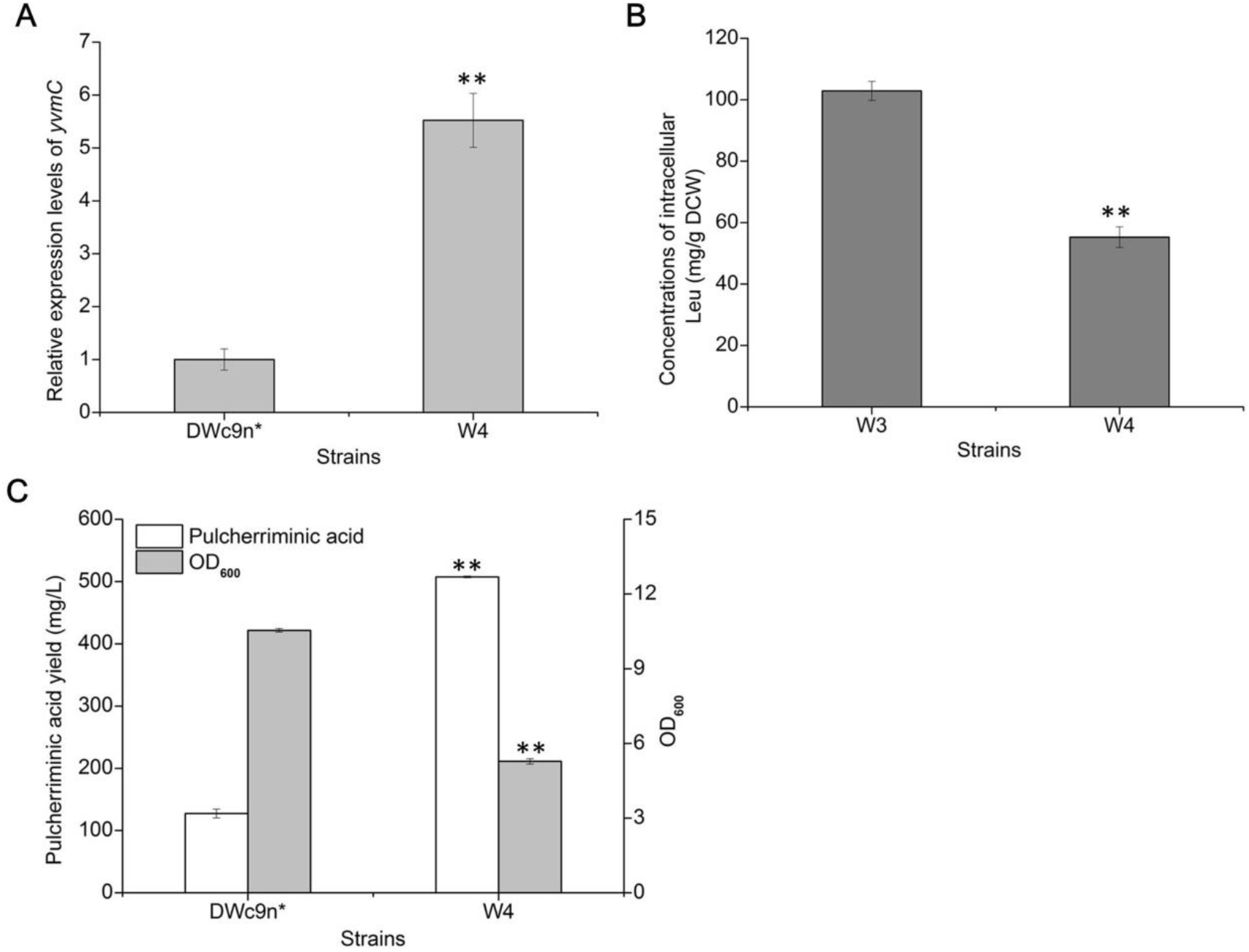
Effect of *yvmC-cypX* overexpression on pulcherriminic acid production. A: Gene transcriptional levels of the *yvmC-cypX* operon at 20 h; B: The concentrations of intracellular Leu at 24 h C: Pulcherriminic acid yield & OD_600_. All samples were performed in three replicates, and data were presented as the mean ± the standard deviation for each sample point. All data were conducted to analyze the variance at P < 0.05 and P < 0.01, and a *t* test was applied to compare the mean values using the software package Statistica 6.0 ∗, P < 0.05 and ∗∗, P < 0.01 indicated the significance levels between recombinant strains and control.

### Overexpression of the transporter gene *yvmA* promoted pulcherriminic acid secretion

The major facilitator superfamily (MFS) like transporter YvmA, which was proved to be responsible for the secretion of pulcherriminic acid (18), was overexpressed through constructing the W4/pHY-*yvmA* strain. The DWc9n*/pHY300 strain was set as a control. In W4/pHY-*yvmA*, the transcriptional level of *yvmA* was 345% higher than in DWc9n*/pHY300 (Fig. 5A), and the concentration of intracellular Leu was 48.7 mg/g DCW at 24 h (Fig. 5B). In addition, our results showed that the intracellular pulcherriminic acid content in W4/pHY-*yvmA* was 8.4 mg/g DCW, which was significantly lower than in W4/pHY300 (31.5 mg/g DCW) (Fig. 5D). At 48 h, the pulcherriminic acid yield of W4/pHY-*yvmA* reached 556.1 mg/L, increased by 335% and 340% compared to the control strain DW2c9n*/pHY300 and DWc9n*, respectively (Fig. 5C). These results showed that the efficiency secretion of pulcherriminic acid would benefit its biosynthesis.

**Fig. 5.**
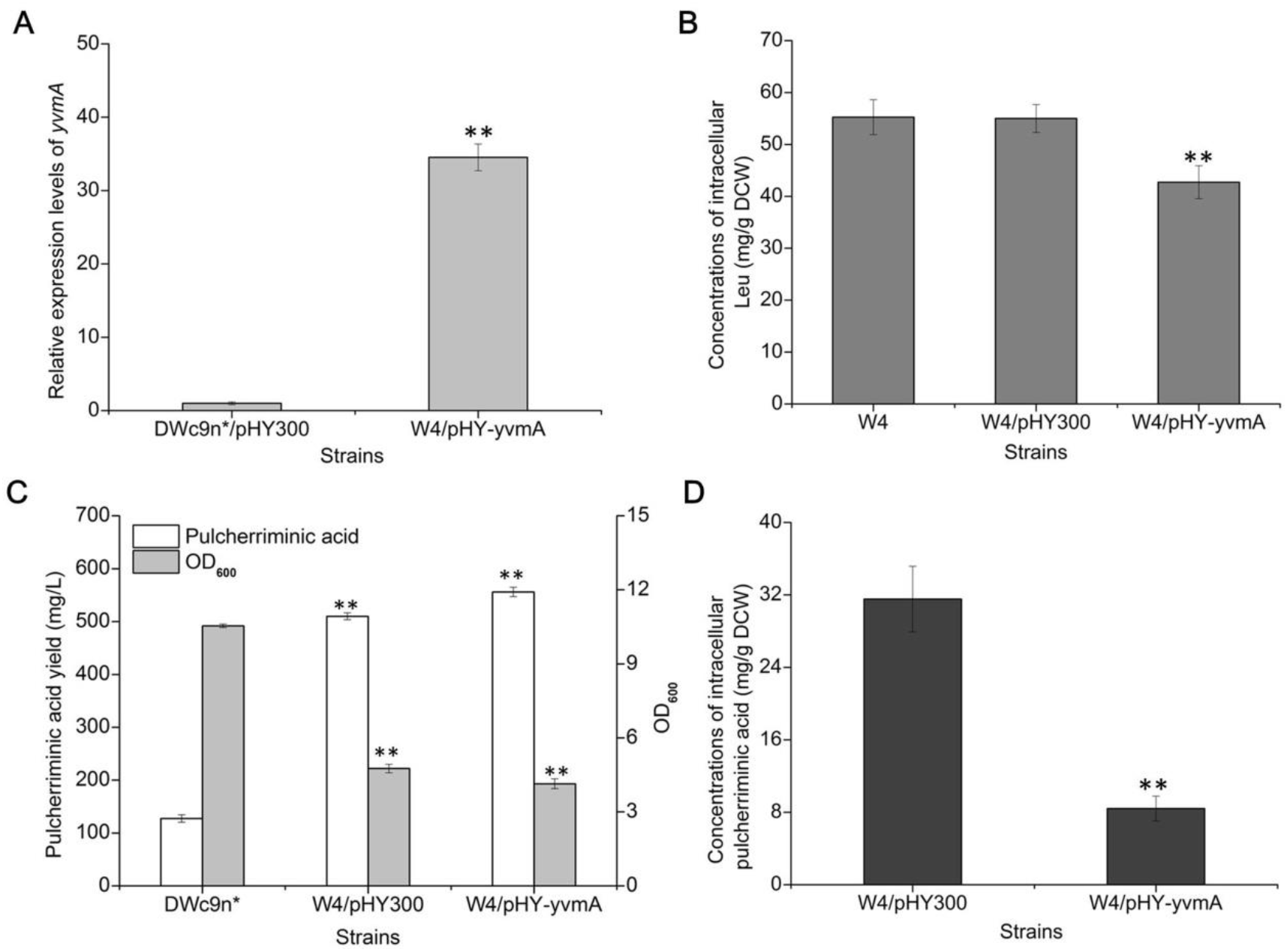
Effect of *yvmA* overexpression on pulcherriminic acid production. A: Gene transcriptional levels of the *yvmA* at 20 h; B: The concentrations of intracellular Leu at 24 h; C: Pulcherriminic acid yield & OD_600_; D: The concentrations of intracellular pulcherriminic acid at 44 h. All samples were performed in three replicates, and data were presented as the mean ± the standard deviation for each sample point. All data were conducted to analyze the variance at P < 0.05 and P < 0.01, and a *t* test was applied to compare the mean values using the software package Statistica 6.0 ∗, P < 0.05 and ∗∗, P < 0.01 indicated the significance levels between recombinant strains and control.

Furthermore, the process curves of *B. licheniformis* DWc9n*/pHY300 and W4/pHY-*yvmA* were measured, and the substrate consumption and byproduct synthesis rates were measured. The cells get the highest density at 44 h, and followed by a minimal decreasing from 44 h to 48 h (Fig. 6A). The growth trend of W4/pHY-*yvmA* was similar with that of DWc9n*/pHY300 from 0 to 36 h, but with a lower growth rate, and the cell density declined after 36 h. Meanwhile, pulcherriminic acid produced by DWc9n*/pHY300 was increased slightly throughout the process, and reached 130.5 mg/L at 48 h. The maximum pulcherriminic acid yield of W4/pHY-*yvmA* reached 556.1 mg/L, increased by 328% compared to DWc9n*/pHY300. In W4/pHY-*yvmA*, the pulcherriminic acid yield per unit cell was 514.8 mg/g DCW increased by 970% as compared to DWc9n*/pHY300 (48.2 mg/g DCW). Moreover, the higher growth rate of DWc9n*/pHY300 resulted in faster glucose consumption in the early growth stage. After 24 h, the pulcherriminic acid levels of W4/pHY-*yvmA* strongly increased and glucose levels in the culture medium dropped (Fig. 6B). In addition, the total concentration of the main byproducts acetoin and 2,3-BD of W4/pHY-*yvmA* was 8.5 g/L, which decreased by 19.1% as compared with that of DWc9n*/pHY300 (10.4 g/L) (Fig. 6C). These results indicating that multistep metabolic engineering strategy strengthening the carbon flux towards the pulcherriminic acid biosynthesis.

**Fig. 6.**
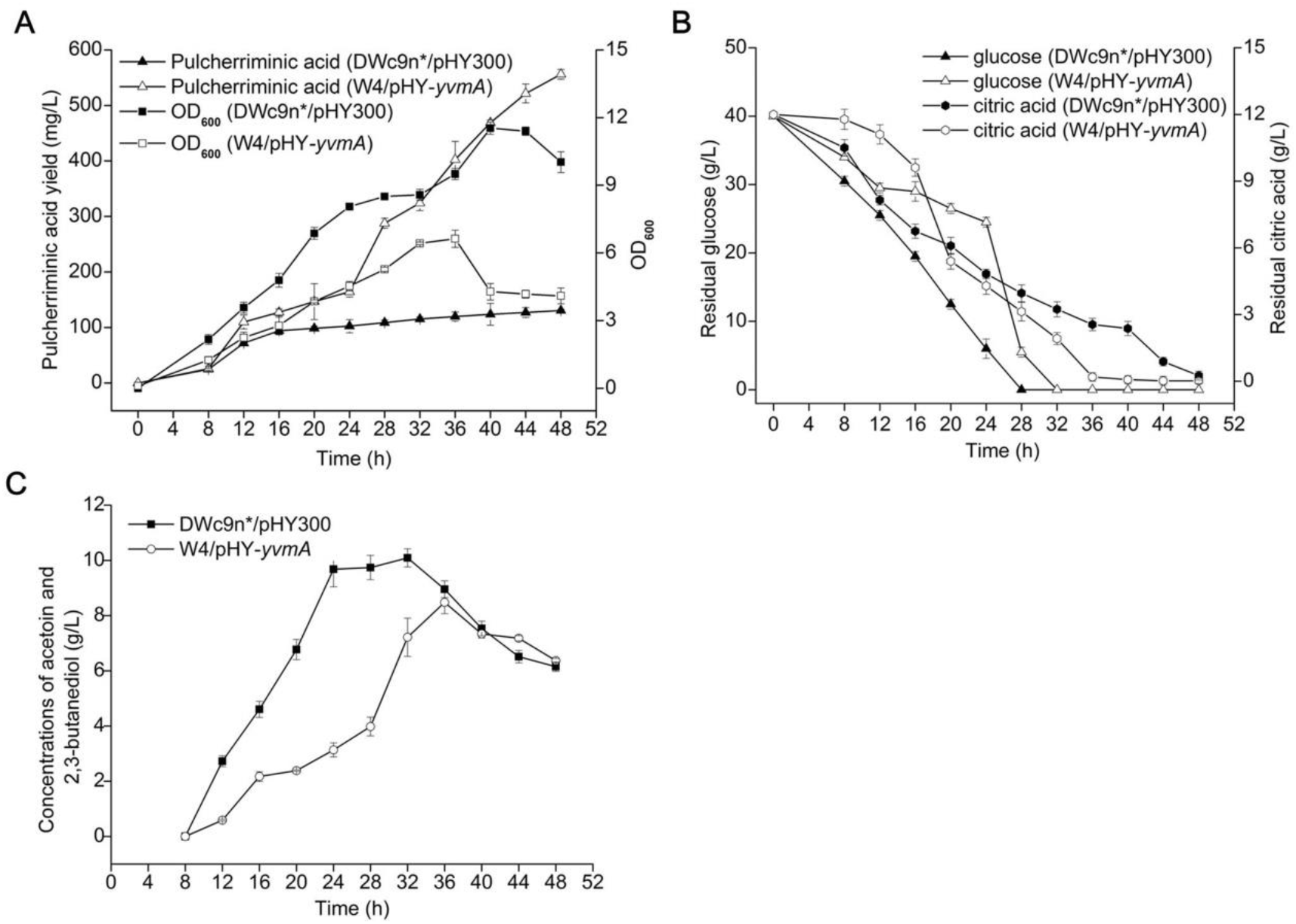
The process curves of *B. licheniformis* DWc9n*/pHY300 and W4/pHY-*yvmA* during pulcherriminic acid production. A: Pulcherriminic acid & OD_600_; B: Residual glucose and citric acid; C: Acetoin and 2,3-butanediol. All samples were performed in three replicates, and data were presented as the mean ± the standard deviation for each sample point.

## Discussion

Pulcherriminic acid may have a wide range of applications as a bacteriostatic agent. (26, 27). Although several studies have focused on the synthesis and metabolic regulation of pulcherriminic acid (28, 29), this is the first metabolic engineering approach for achieving high-level pulcherriminic acid production. In this study, the pulcherriminic acid synthetic capability was significantly increased by rewiring the metabolic pathway of *B. licheniformis*, and the strain W4/pHY-*yvmA* with a 340% increased pulcherriminic acid yield was generated.

Precursor supply plays the critical role in metabolite biosynthesis (30). Here, Leu, derived from pyruvate, acted as the initial precursor for pulcherriminic acid synthesis. Since the low concentration of intracellular Leu in *B. licheniformis* DW2 (31), Leu supply might be a bottleneck for pulcherriminic acid production. Vogt et al. (2014) have obtained a valid L-Leu production strain of *Corynebacterium glutamicum* by deleting the feedback-resistent and the repressor LtbR for L-Leu synthesis, leading to a Leu yield increase to 24 g/L under optimized fed-batch fermentation (32). In addition, Zhu et al. (2018) have improved the accumulations of intracellular BCAAs by overexpressing the BCAA importer BrnQ, which led to a 22.4% increase of bacitracin yield (31). In the present work, the concentration of intracellular Leu was increased by 230% via overexpressing *ilvBHC-leuABCD* and deleting *bkdAB*, which led to a 49% increase of pulcherriminic acid production. Meanwhile, overexpressing aminoacyl tRNA synthetase (LeuS) was also served as the efficient strategy for metabolite synthesis. Xia et al. (2010) strengthen the expression of glycyl-tRNA synthetase to enhance the glycyl-tRNA pool, which efficiently produced a specific protein from the spider *Nephila clavipes* with a molecular weight of 284.9 kDa (33). Here, the aminoacyl tRNA synthetase *leuS* was overexpressed to increase the formation of leucyl-tRNA, which led to a 190% increase of pulcherriminic acid production and a decrease of intracellular Leu by 43%. In short, our results show that precursor supply played a vital role in pulcherriminic acid synthesis and pulcherriminic acid yield could be increased via strengthening the Leu and leucyl-tRNA supplies.

The CDPS proteins recognize the aa-tRNA in an RNA sequence-independent manner, which resulted in the lower substrate specificity (25). Therefore, several kinds of other cyclodipeptides not only cyclo(L-Leu-L-leu) might be synthesized as by-products simultaneously, which could limit the synthesis efficiency of target cyclo-amino acids and metabolites (34). Although most CDPS show a degree of substrate promiscuity, they mainly incorporate five hydrophobic amino acids (Phe, Leu, Tyr, Met and Trp) into cyclodipeptides. For instance, the cyclodipeptides catalyzed by CDPS AlbC from *Streptomyces sinensis* showed at least 12 different amino acid combinations, such as cyclo(L-Phe-L-Leu), cyclo(L-Phe-L-Tyr), cyclo(L-Phe-L-Phe), cyclo(L-Phe-L-Met) or cyclo(L-Tyr-L-Met) (Sauguet et al., 2011). Six cyclodipeptides, including cyclo(L-Tyr-L-Tyr), cyclo(L-Tyr-L-Phe), cyclo(L-Tyr-L-Leu), cyclo(L-Tyr-L-Ala), cyclo(L-Tyr-L-Met) and cyclo(L-Tyr-L-Trp), were synthesized by CDPS Rv2275 of *Mycobacterium tuberculosis* (35). Based on previous research, CDPS YvmC from *Bacillus* also showed a low substrate specificity and approximately 40% of the products were cyclo(L-Leu-L-Phe) and cyclo(L-Leu-L-Met), but not the target product cLL (15). Our results show that the concentration of intracellular Phe was significantly decreased in *yvmC-cypX* overexpression strain (**Fig. S1**), synchronously with that of intracellular Leu, which suggested that cyclo(L-Leu-L-Phe) might also be accumulated in strain W4. Unfortunately, the concentration of intracellular Met has not been detected in this research. Despite this, the catalysis specificity of YvmC should be engineered to improve pulcherriminic acid production in our subsequent work.

Unlike the antibiotics producing strains, which get self-resistant by secreting its products, the growth of the pulcherriminic acid producing strain was also repressed by iron starvation (18). Moreover, the synthesis of cyclic dipeptides is rigorously regulated in microorganisms (19). Based on our previous research, the synthesis of pulcherriminic acid is regulated by the transcriptional regulators AbrB, YvnA and YvmB, to reach iron homeostasis under a range of iron stress (18). Similar dual-regulation systems were also confirmed in the biosynthesis of several other secondary metabolites (17, 36). Gondry et al. (2009) have measured the catalytic efficiencies of eight CDPSs, such as YvmC and AlbC, and their results showed yields of cyclic dipeptides at 16.6 and 117.5 mg/L (25). In this study, the strategy of promoter replacement might have broken the rigorous regulatory system of *B. licheniformis* and allowed a pulcherriminic acid yield of 556.1 mg/L in the strain W4/pHY-*yvmA* the highest yield of pulcherriminic acid reported so far.

**Fig. 7.**
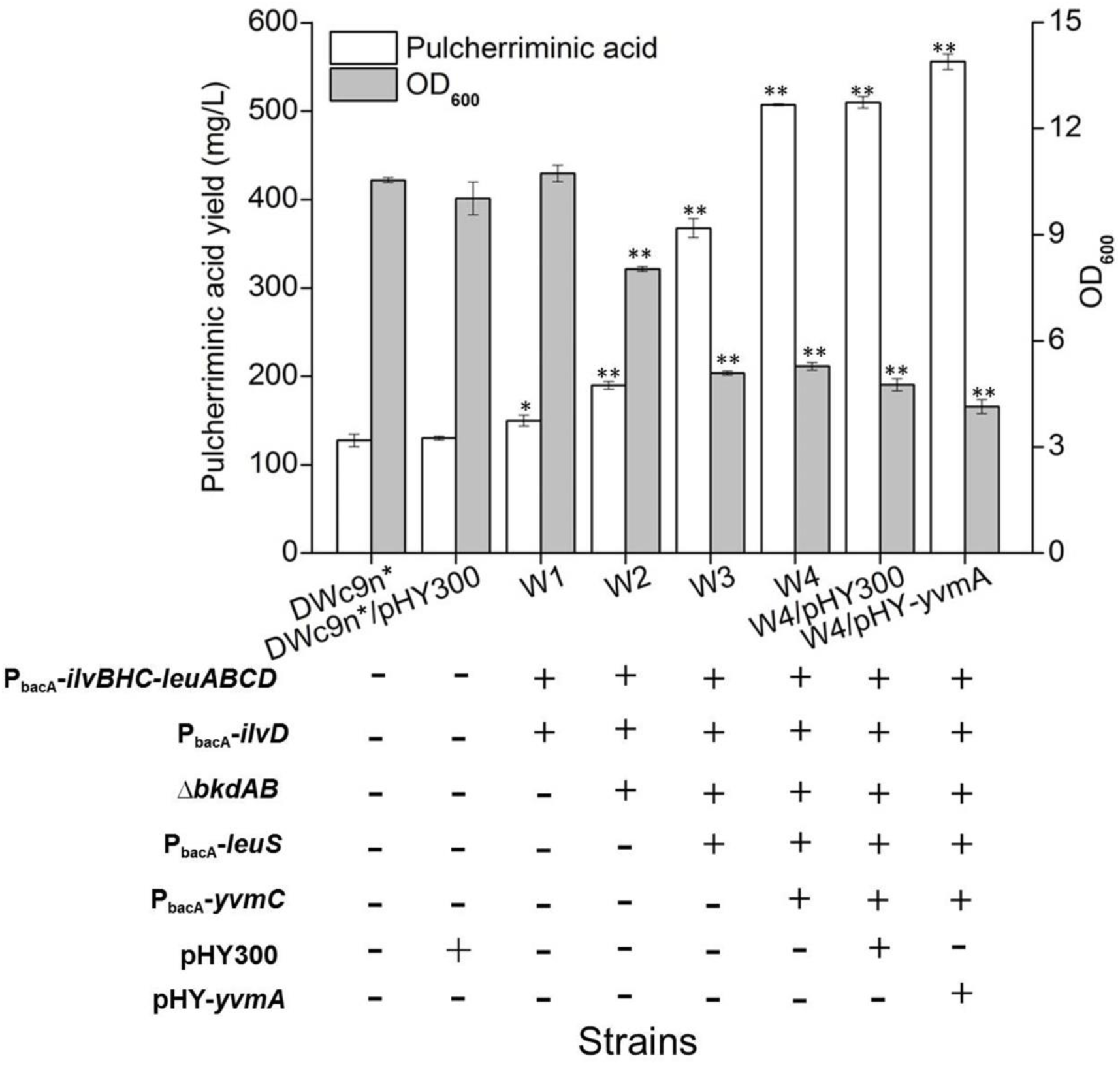
The relationship between pulcherriminic acid yields and OD_600_ among these recombinant strains. All samples were performed in three replicates, and data were presented as the mean ± the standard deviation for each sample point. All data were conducted to analyze the variance at P < 0.05 and P < 0.01, and a *t* test was applied to compare the mean values using the software package Statistica 6.0 ∗, P < 0.05 and ∗∗, P < 0.01 indicated the significance levels between recombinant strains and control.

## Conclusion

In this study, Leu and leucly-tRNA supply, cyclodipeptide synthetase and pulcherriminic acid transporter were increased in strain to maximise pulcherriminic acid production. The best performing strain W4/pHY-*yvmA* had a maximum yield of 556.1 mg/L of pulcherriminic acid, which represented a 340% increase as compared to the original strain DWc9n*. To the best of our knowledge, this is the highest yield of pulcherriminic acid reported so far. Collectively, this study provided an efficient strategy for enhancing the production of pulcherriminic acid, that could also be applied for strain improvement and high-level production of other cyclodipeptides.

## Author’s contribution

D Wang and S Chen designed the study. S Wang, D Wang, X Li and D Zhang carried out the molecular biology studies and constructing engineering strains. S Wang, H Wang, D Zhang and J Zhang carried out the fermentation studies. S Wang, D Wang, Y Zhan, D Cai, X Ma and S Chen analyzed the data and wrote the manuscript. All authors read and approved the final manuscript.

## Acknowledgments

This work was supported by the National Key Research and Development Program of China (2018YFA0900303, 2015CB150505), the Technical Innovation Special Fund of Hubei Province (2018ACA149).

## Supporting Information

All the primers sequences for strain construction and qRT-PCR were listed in **Table S1**. The concentrations of intracellular Phe among those recombinant strains at 24 h were provided in **Fig. S1**.

